# A single-cell anatomical blueprint for intracortical information transfer from primary visual cortex

**DOI:** 10.1101/148031

**Authors:** Yunyun Han, Justus M Kebschull, Robert AA Campbell, Devon Cowan, Fabia Imhof, Anthony M Zador, Thomas D Mrsic-Flogel

**Affiliations:** Biozentrum, University of Basel, 4056 Basel, Switzerland; Department of Neurobiology School of Basic Medicine and Tongji Medical College, Huazhong University of Science and Technology, Wuhan, China; The Institute for Brain Research, Collaborative Innovation Center for Brain Science, Huazhong University of Science and Technology, Wuhan, China; Watson School of Biological Sciences, Cold Spring Harbor, NY 11724, USA; Cold Spring Harbor Laboratory, Cold Spring Harbor, NY 11724, USA; Sainsbury Wellcome Centre, University College London, London, UK

## Abstract

The wiring diagram of the neocortex determines how information is processed across dozens of cortical areas. Each area communicates with multiple others via extensive long-range axonal projections ^1–6^, but the logic of inter-area information transfer is unresolved. Specifically, the extent to which individual neurons send dedicated projections to single cortical targets or distribute their signals across multiple areas remains unclear^5,7–20^. Distinguishing between these possibilities has been challenging because axonal projections of only a few individual neurons have been reconstructed. Here we map the projection patterns of axonal arbors from 591 individual neurons in mouse primary visual cortex (V1) using two complementary methods: whole-brain fluorescence-based axonal tracing^21,22^ and high-throughput DNA sequencing of genetically barcoded neurons (MAPseq)^23^. Although our results confirm the existence of dedicated projections to certain cortical areas, we find these are the exception, and that the majority of V1 neurons broadcast information to multiple cortical targets. Furthermore, broadcasting cells do not project to all targets randomly, but rather comprise subpopulations that either avoid or preferentially innervate specific subsets of cortical areas. Our data argue against a model of dedicated lines of intracortical information transfer via “one neuron – one target area” mapping. Instead, long-range communication between a sensory cortical area and its targets may be based on a principle whereby individual neurons copy information to, and potentially coordinate activity across, specific subsets of cortical areas.

---

The function of a neuron can be defined both by its receptive field^24^, which describes the inputs that drive it to fire, and its projective field^25^, which describes its impact on other neurons. The projective field of a neuron is determined by the pattern of its axonal projection, and thereby governs how information propagates between brain areas. Anatomical studies, largely based on retrograde tracing methods, suggest an abundance of intracortical projection neurons in sensory neocortex whose axons appear to innervate single target areas^5,7–9^, raising the possibility that information is distributed via ensembles of dedicated pathways that are functionally tailored to each target^10–16^. For example, neurons in mouse primary visual cortex (V1) that innervate the posteromedial (PM) or anterolateral (AL) area appear to match the spatial and temporal frequency preference of these target areas^10,26,27^. Similarly, neurons in the primary somatosensory cortex projecting to either primary motor cortex or the secondary somatosensory area comprise largely non-overlapping populations with distinct physiological and functional properties^12–14^. These findings suggest that dedicated lines — specialized subpopulations of neurons that preferentially target a single downstream area (Fig. 1a, top) — may represent a fundamental mode of cortico-cortical communication. Alternatively, sensory cortical neurons might broadcast to multiple targets^5,9,17–21^, either randomly (Fig. 1a, middle), or by targeting specific sets of areas (Fig. 1a, bottom). These three models of cortical architecture have different implications for inter-areal communication underlying sensory processing in hierarchical networks. We therefore set out to distinguish among them, using anterograde anatomical methods to map the long-range axonal projection patterns of individual neurons in mouse primary visual cortex (V1), an area that distributes visual information to multiple cortical and subcortical targets^1,2,6^.

**Figure 1:**
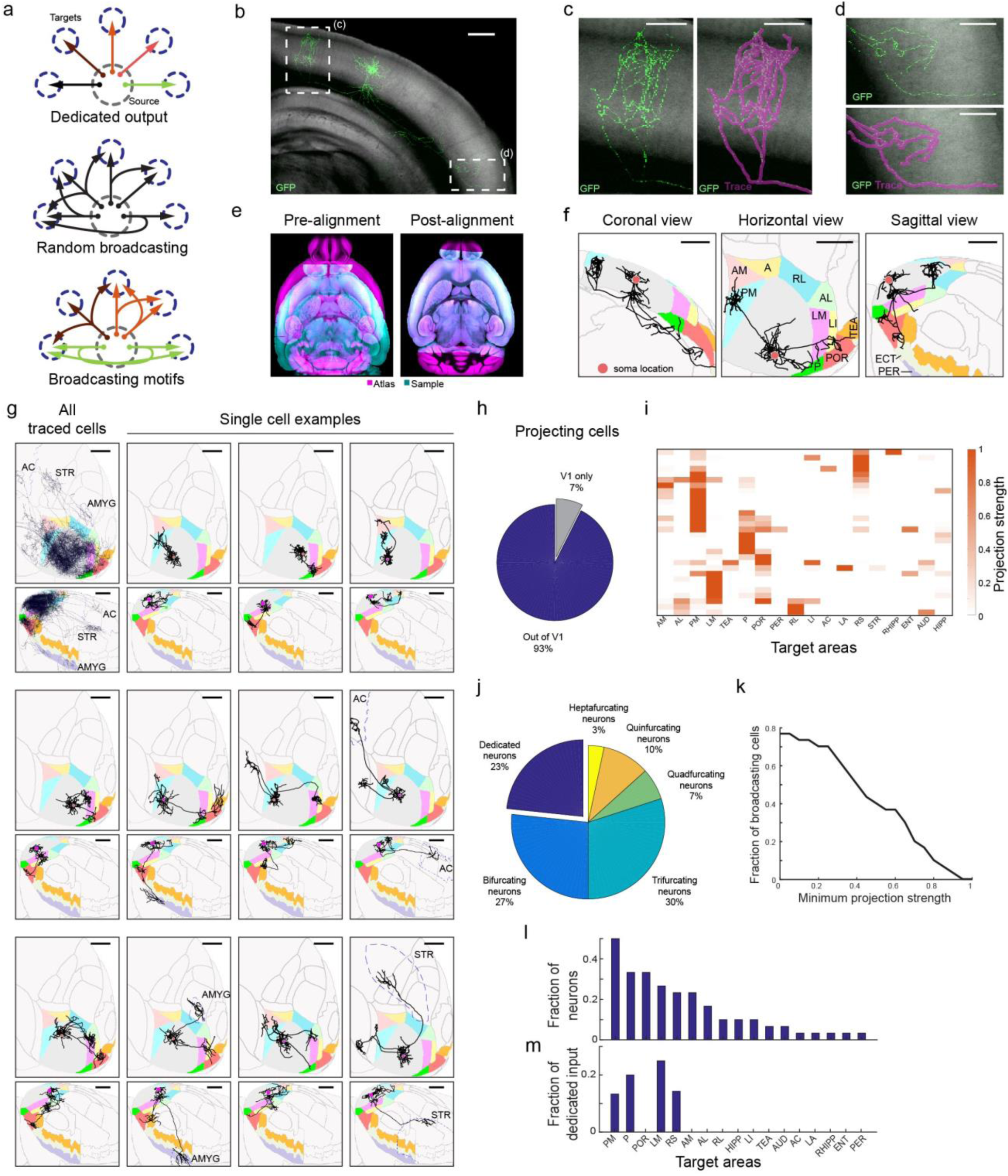
Brain-wide single-cell tracing reveals the diversity of axonal projection patterns of layer 2/3 V1 neurons, with most cells projecting to more than one target area. (**a**) Three hypothetical modes of inter-areal information transfer from one area to its multiple targets. Neurons (arrows) could each project to a single area (top) or to several areas either randomly (middle) or in predefined projection patterns (bottom). (**b**) Maximum projection of an example GFP-filled neuron coronal view acquired by serial-section 2-photon microscopy. Auto-fluorescence from the red channel is used to show the brain’s ultrastructure (gray background). Scale bar = 600 μm. (**c**-**d**) Higher magnification of the medial (**c**) and lateral (**d**) axonal arbor of the example cell. Scale bar = 300 μm. (**e**) Horizontal section through a sample brain (cyan) and Allen reference atlas (ARA; magenta) before (left) and after (right) rigid and non-rigid transformation of the brain to the atlas. (**f**) Coronal, sagittal and horizontal projections of the traced example cell overlaid in ARA space. Target cortical areas are coloured as indicated. Areas: A, anterior; AL: anterolateral; AM: anteromedial; LI: lateroitermediate; LM: lateral; P: posterior; PM: posteromedial; POR: postrhinal; RL: rostrolateral; TEA: temporal association; ECT: ectorhinal; PER: perirhinal. Scale bar = 1 mm. (**g**) Overlay of all traced single neurons (top left) and 11 example cells in Allen Reference Atlas (ARA) space; horizontal view (upper panel) and sagittal view (lower panel). Dashed outlines label non-visual target areas: AC: anterior cingulate cortex; STR, striatum; AMYG: amygdala. Note that these images are for illustration purposes only because a 2D projection cannot faithfully capture the true axonal arborisation pattern in 3D. Scale bar = 1 mm. (**h**) Pie chart illustrating the fraction of traced single neurons that project to at least one target area outside V1, where at least 1 mm of axonal innervation is required for an area to be considered a target. (**i**) Projection pattern of all GFP-filled V1 neurons targeted randomly (upper panel, n=31). The colour-code reflects the projection strengths of each neuron, determined as axon length per target area, normalized to the axon length in the target area receiving the densest innervation. Only brain areas that receive input form at least one neuron, as well as striatum, are shown. Areas: AUD: auditory cortex; ENT: entorhinal; HIPP: hippocampus; LA: lateral amygdala; RHIPP: retrohippocampal region; RS: retrosplenial. (**j**) The number of projection targets for every neuron that projects out of V1. (**k**) The proportion of cells targeting more than one area, when projection targets that receive projections weaker than the indicated projection strength are ignored. For each neuron, projection strengths are normalized to axon length in the target area receiving the densest innervation. (**l**) The fraction of neurons projecting to each of the 18 target areas of V1. (**m**) The fraction of neurons innervating a single target area (‘dedicated’ projection neurons) out of all neurons that innervate that area.

We used single-cell electroporation of a GFP-encoding plasmid to label up to six layer 2/3 cells in the right visual cortex of each mouse. After allowing 3-10 days for GFP expression we imaged the axonal projections of the labeled neurons by whole-brain serial two-photon tomography with 1x1x10 μm resolution^28,29^ (Fig. 1b). We then traced each fluorescently-labeled cell (Fig. 1c,d; n = 71) and registered each brain to the Common Coordinate Framework 3 of the Allen Reference Atlas (Allen Institute, 2015) in order to segment and identify the brain areas in which axonal terminations were observed (Fig. 1e,f). To assess the extent of axonal labelling with GFP, we electroporated neurons labelled retrogradely from the ipsilateral striatum — a distal projection target of V1 — and in all cases observed axonal terminations therein (n = 9/9 cells; Extended Data Fig. 1). Nonetheless, a fraction of reconstructed V1 neurons contained axon collaterals beyond V1 that terminated abruptly without arborizing (n = 28; Extended Data Fig. 2). These neurons were excluded from further analysis because it is impossible to ascertain whether such abrupt terminations are real or due to incomplete axonal filling. Note that we did not exclude neurons with abrupt terminations of contralaterally projecting branches (compare ref^14^), instead restricting our analysis to ipsilaterally-projecting axons.

We analysed the ipsilateral projection patterns of 38 pyramidal neurons in layer 2/3, including 31 neurons in area V1 (Fig. 1g, Extended Data Fig. 3 and 4) and 7 neurons in nearby higher visual areas (Extended Data Fig. 5). Inspection of individual axonal arbors of V1 neurons revealed a high degree of projectional diversity among neurons (Fig. 1g, Extended Data Fig. 3 and 4) that is obscured in bulk projection data^1,2^ (Fig. 1g, top left).

Almost all layer 2/3 cells (93%) projected out of V1 (Fig. 1h) to one or more of 18 target areas (including striatum; Extended Data Fig. 1) in the telencephalon (Fig. 1i). To mitigate errors arising both from technical noise in atlas registration and from subject-to-subject variability in the boundaries between brain areas, we excluded low-confidence “buffer zones” of 100 μm around the area boundaries from analysis, and define as a “target” only those areas that received over 1 mm of axonal input from an individual cell (see Methods). Eighty-five percent of all projection patterns appeared only once, highlighting the diversity of long-range projection fields of V1 neurons.

The majority (77%) of reconstructed layer 2/3 projection neurons sent axons to more than one target area, with some targeting up to seven areas throughout the brain (Fig. 1j). While individual neurons innervated different target areas with different axonal densities, and thus might influence the computations in one area more than another, we found that a large fraction of broadcasting cells innervated more than one target with comparable strengths (Fig. 1k). Posteromedial visual (PM) area was the most common target, followed by posterolateral (P), postrhinal (POR) and lateromedial (LM) areas (Fig. 1l). Even when the analysis was restricted to neurons that projected to at least one of six nearby cortical visual areas (LI, LM, AL, PM, AM, RL), we found that half projected to two or more of these areas (Extended Data Fig. 6a-e). The fraction of input provided by dedicated projection neurons to any area comprised no more than 25% of the total (Fig. 1m), and most target areas received no dedicated input. These conclusions were robust to changes in the size of the border exclusion zone between neighbouring areas and the minimum projection strength in the target area (Extended Data Fig. 6f-h). Similar to projections from V1, all seven reconstructed neurons whose cell bodies resided in nearby higher visual areas also projected to more than one target area (Extended Data Fig. 5). Our results thus reveal that most layer 2/3 neurons distribute information to multiple areas, rather than projecting to single targets.

We next investigated whether broadcasting cells targeted cortical areas at random, or whether there was higher-order structure, i.e. whether subpopulations of neurons preferentially target or avoid specific subsets of areas. Here we define higher-order structure in terms of the connection patterns predicted by the per-neuron (first order) probability of projecting to each target. For example, if the probability of any given neuron projecting to area A is 0.5 and the probability of projecting to area B is also 0.5 then we would expect P(A∩B)=P(A)*P(B)=0.25 of all neurons to project to both A and B. Significant deviations from this expectation would indicate organization into non-random projection motifs. Probing for high order structure requires large datasets, because, if a sample size of *N* neurons is required to estimate the first order probabilities, then a sample size of *N^2^* is needed to estimate pairwise probabilities with comparable accuracy. Although single neuron reconstruction provides unrivaled spatial resolution, despite increases in throughput for data acquisition^21,22^, the tracing of axons remains labor intensive.

We therefore used a higher throughput method, MAPseq^23^, to obtain a large number of single neuron projections for higher-order statistical analysis. In a MAPseq experiment, hundreds or thousands of neurons are labeled uniquely with random RNA sequences (barcodes) by a single injection of a library of barcoded Sindbis virus (Supplemental Note 1). The barcodes are expressed and then actively transported into the axonal processes of each labeled neuron, where they can be read out by high throughput barcode sequencing after dissection of potential target areas. The abundance of each barcode sequence in each area serves as a measure of the projection strength of the corresponding barcode-labeled neuron. MAPseq thus simultaneously maps the projections of all labeled neurons to dissected target areas, and therefore allows in-depth analysis of projections to a smaller set of targets.

We used MAPseq to map the projection patterns of 553 neurons from V1 to six higher visual areas — LI, LM, AL, AM, PM and RL — that can be identified reliably by intrinsic signal imaging *in vivo* and dissected *ex vivo* for barcode sequencing (Fig. 2a,b; Extended Data Fig. 7; see Methods). To avoid virus spillover from V1 into adjacent areas, we made focal injections of the MAPseq virus to yield 100-200 traced cells per animal. Consistent with the analysis of fluorescence-based single neuron reconstructions restricted to the six higher visual areas (Fig. 2c, left), almost half (44%) of all MAPseq neurons projected to more than one area (Fig. 2c, right). Furthermore, the projection patterns obtained by fluorescence-based tracing were statistically indistinguishable from those obtained by MAPseq (using a bootstrap procedure; see Supplemental Note 2), whereas neurons with projection strengths sampled from a uniform distribution were markedly different (Fig. 2d). Thus the findings from the MAPseq dataset were consistent with those from single neuron tracing.

**Figure 2:**
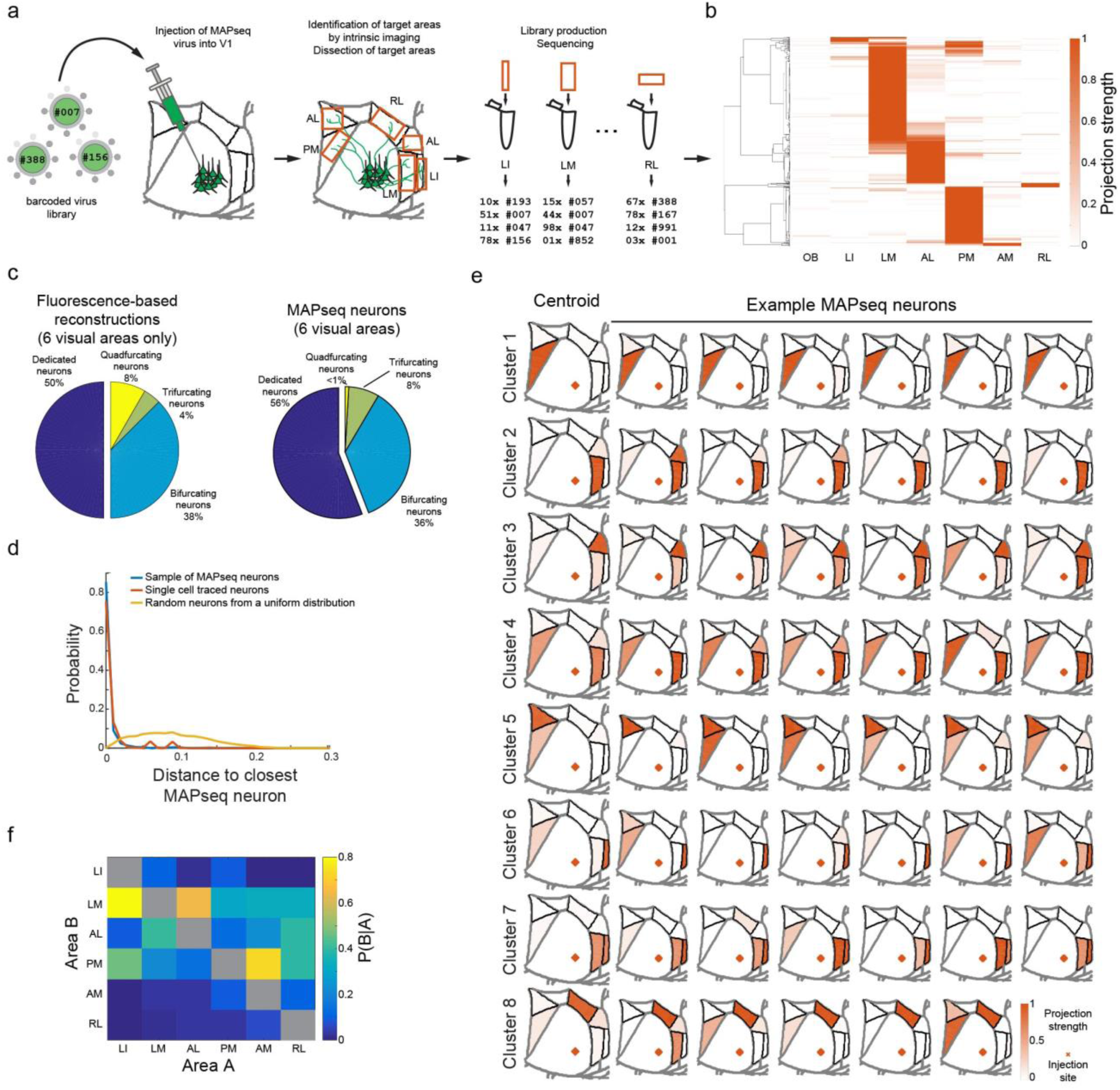
MAPseq projection mapping reveals a diversity of projection motifs. (**a**) Overview of the MAPseq procedure. Six target areas were chosen for analysis: LI, LM, AL, PM, AM and RL. (**b**) Projection strength in the six target areas, as well as the olfactory bulb (OB) as a negative control, of 553 MAPseq-mapped neurons. Projection strengths per neuron are defined as the number of barcode copies per area, normalized to the efficiency of sequencing library generation and to the neuron’s maximum projection strength (n=4 mice). (**c**) Number of projection targets of V1 neurons when considering the six target areas only, based on the fluorescence-based axonal reconstructions (left) or the MAPseq data (right). (**d)** Distribution of cosine distances obtained by a bootstrapping procedure between MAPseq neurons (blue), fluorescence-based single neuron reconstructions and MAPseq neurons (orange), or random neurons (with projection strengths sampled from a uniform distribution) and MAPseq neurons (yellow). The distance distributions obtained from MAPseq neurons and fluorescence-based single-neuron reconstructions are statistically indistinguishable (Kolmogorov-Smirnov two sample test; p=0.94), whereas the distributions obtained from both MAPseq neurons or fluorescence-based reconstructed neurons are statistically different form the distribution obtained using random neurons (Kolmogorov-Smirnov two sample test; p<10^-3^). (**e**) Centroids and example cells for eight clusters obtained by k-means clustering of all MAPseq cells using a cosine distance metric. Target areas are coloured to indicate the projection strength of the plotted neuron. Projections strengths are normalize as in (**b**). (**f**) The probability of projecting to one area (Area A) given that the same neuron is projecting to another area (Area B) based on the MAPseq dataset.

We first catalogued the diversity of single neuron projection patterns from V1 to six higher visual areas by unsupervised clustering of the MAPseq dataset (k-means clustering with a cosine distance metric). These projectional data were best described by eight clusters (Fig. 2e, Extended Data Fig. 8), of which all but one contained cells targeting more than one area. The most common partners for broadcasting neurons were LM and PM, consistent with the fact that a large fraction of neurons targeted these areas (Fig. 2f).To uncover the existence of non-random projection motifs in the MAPseq dataset, we measured the likelihood of specific bi-, tri- or quadfurcations, by comparing them to expected probabilities of these divergent projections (assuming independence between each projection type; Fig. 3a,b). This analysis identified six projection motifs that were significantly over- or under-represented after a correction for multiple comparisons (Bonferroni adjustment; Fig. 3b,c). Together, these six projection motifs represented 73% of all broadcasting cells identified by MAPseq. Therefore the majority of V1 cells projecting to multiple target areas do so in a non-random manner, suggesting that broadcasting motifs reflect several sub-classes of projection neurons for divergent information transfer from V1 to higher visual areas.

**Figure 3:**
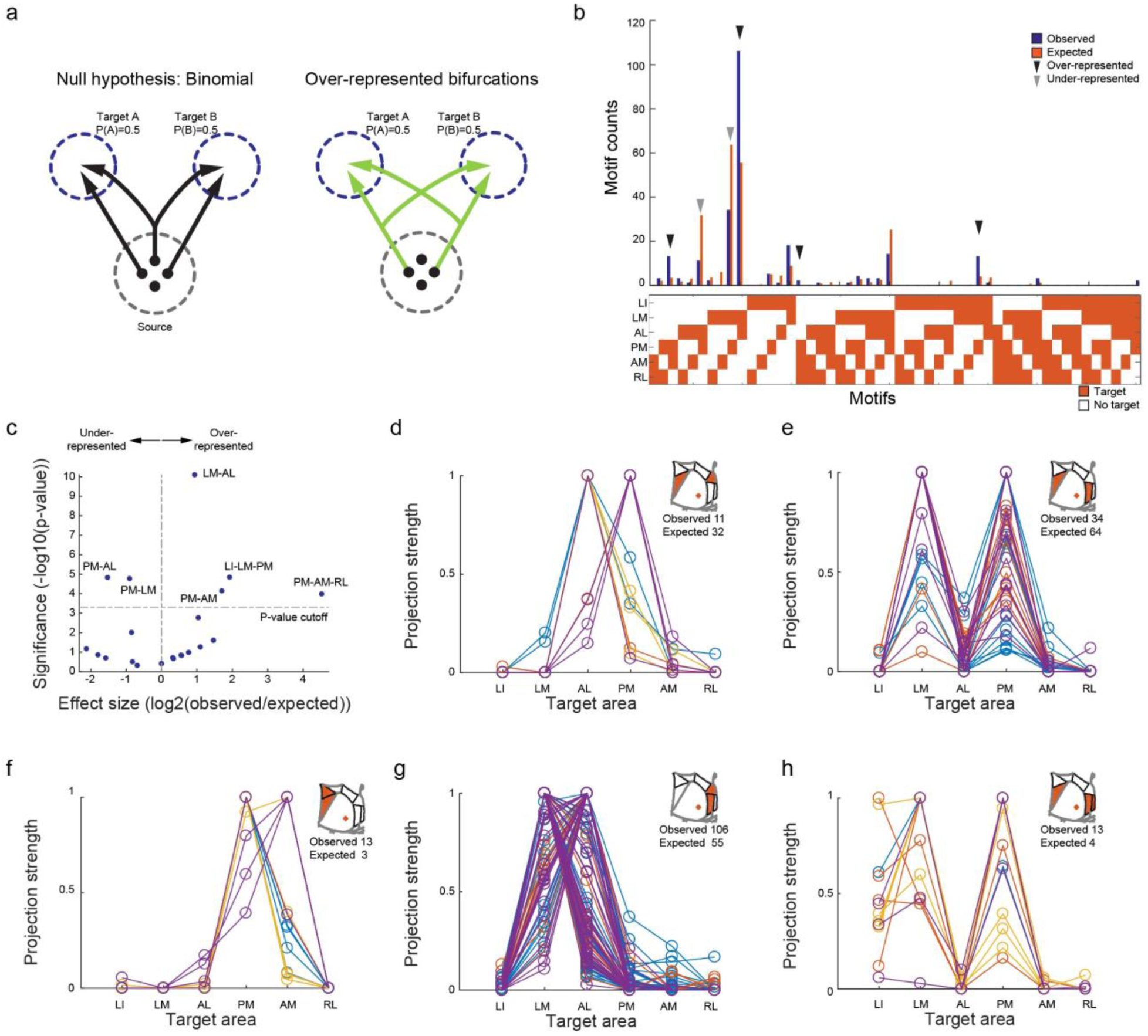
Over- and under-represented projection motifs of neurons in primary visual cortex. (**a**) The null hypothesis of independent projections to two target areas (left) and an example deviation (over-represented bifurcation) from the null hypothesis (right). (**b**) The observed and expected abundance of all possible bi-, tri- and quadfrucation motifs in the MAPseq dataset. Significantly over- or under-represented motifs are indicated by black and grey arrowheads, respectively. (**c**) Statistical significance of over- and under-represented broadcasting motifs and associated effect sizes. (**d**-**h**) The projection strengths of the individual neurons (one per line) giving rise to the six under-represented (**d**,**e**) or over-represented (**f**-**h**) projection motifs. For each neuron, the projections strength in each target area is normalized to the neuron’s maximum projection strength. Lines of the same color represent neurons mapped in the same brain (n=4 mice).

The most under-represented broadcasting motif was the bifurcation between areas PM and AL (Fig. 3d). These two areas exhibit distinct visual response properties^26,27^ and receive functionally specialized input from V1^10^, consistent with idea of dedicated projections from V1 into these areas. Moreover, the under-represented population of neurons that do project to both PM and AL was further split into two groups according to projection strength, one that primarily innervates PM and another that primarily innervates AL (Fig. 3d). A second under-represented motif is the bifurcation between PM and LM (Fig. 3e). In contrast to the PM-AL bifurcation, however, the detected PM-LM projecting neurons do not separate cleanly into two classes. Our findings therefore provide an anatomical substrate for the functional dichotomy of areas AL and PM, and suggest that a few ‘dedicated’ output channels can co-exist with a preponderance of broadcasting cells co-innervating multiple targets

In addition to the two under-represented motifs, we also identified four over-represented motifs, i.e. combinations of target areas receiving more shared input from individual V1 neurons than expected from first-order projection statistics (Fig. 3f-h). We found that cells jointly innervating PM and AM were significantly more abundant than expected by chance (Fig. 3f). Resolving the projection strengths within this motif revealed two subpopulations of neurons, one innervating PM more than AM, the other innervating the two areas with similar strength. Moreover, neurons bifurcating to LM and AL were also highly over-represented (Fig. 3g) and comprised the most abundant class of broadcasting cells (Fig. 3b). The most significantly over-represented trifurcation motif was the projection to PM, LM and LI, comprising a relatively homogenous population that projects to LM and PM with similar strengths while slightly less to LI (Fig. 3h). Finally, we discovered the over-representation of the PM-AM-RL trifurcation, but it appeared only rarely in our dataset (Fig. 3b).

These projectional data have implications for the categorization of higher visual areas into putative streams of visual processing in mouse neocortex. Areas AL and PM on the one hand, and LM and LI on the other, have been suggested to belong to dorsal and ventral processing streams in the mouse visual system, respectively^30,31^. Our data indicate that such a distinction is unlikely to originate as a result of segregated V1 input into these areas because they receive a highly degree of shared input (e.g. LM-PM bifurcation, AL-LM bifurcation and PM-LM-LI trifurcation).

In summary, our results provide an insight into the logic by which single neurons in one cortical area distribute information to downstream target areas. Almost all layer 2/3 pyramidal cells projected outside of V1, indicating that V1 neurons concurrently engage in local and distal computations. We found that the single neuron projections beyond V1 were highly diverse, innervating up to seven targets, predominantly in specific, non-random combinations (Fig. 4). These results suggest a functional specialization of subpopulations of projection cells beyond ‘one neuron – one target area’ mapping.

**Figure 4:**
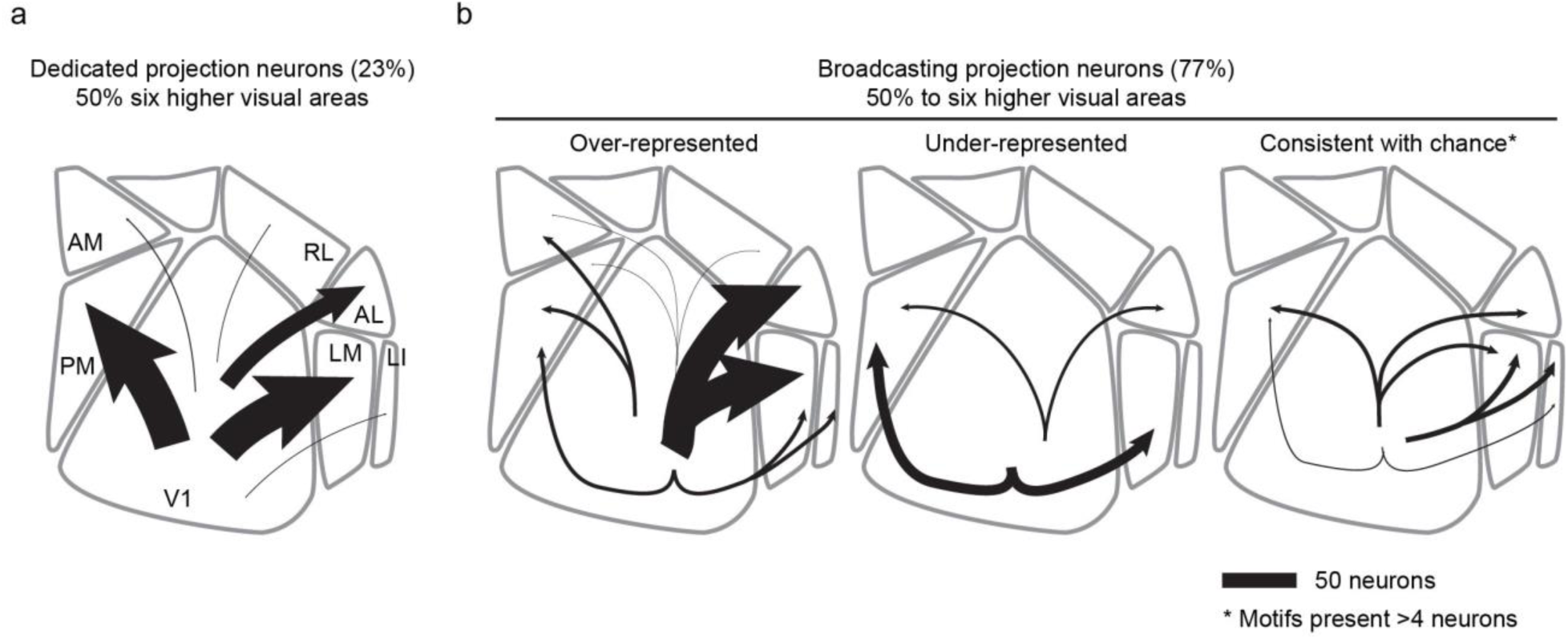
Summary of single-neurons projections from V1. (**a**) Cells targeting single higher visual areas (dedicated projection neurons) comprise the minority of layer 2/3 V1 projection neurons. Among the areas analysed by MAPseq, dedicated projection neurons predominantly innervate cortical areas LM or PM. (**b**) Cells projecting to two or more areas (broadcasting projection neurons) are the dominant mode of information transfer from V1 to higher visual areas. In the six areas analysed by MAPseq, broadcasting neurons innervate combinations of target areas in a non-random manner, including those that are more or less abundant than expected by chance. Line width indicates the absolute abundance of each projection type as observed in the MAPseq dataset.

The fraction of neurons in V1 that broadcast information to multiple targets is considerably greater than previously documented using retrograde methods^7,9,18^. This difference is unlikely caused by differences in the sensitivity with which these approaches detect the projections patterns of individual cells. Instead, anterograde tracing maps projections to many or all targets simultaneously, whereas retrograde tracing typically probes only two or three potential target sites at a time. Because the fraction of neurons projecting to any pair of targets selected for retrograde tracing is relatively low (typically <10%), most neurons will not be doubly labeled in any given experiment; only by sampling many potential targets in a single experiment can the true prevalence of broadcasting be uncovered. Indeed, if we simulate double retrograde tracing based on our MAPseq results, the fractions of bifurcating neurons are comparable to those observed using retrograde methods in primates^7,9,18,19^ (Extended Data Table 1).

We speculate that dedicated projection neurons — which comprise the minority of neurons in V1 — convey visual information tailored to their target area, as suggested previously^10–15^. Indeed, the most under-represented projection motif from V1, the PM-AL bifurcation, innervates two target areas with distinct preferences for visual features^26,27^. In contrast, we suggest that the majority of cells encode information that is shared and in a form suitable for generating visual representations or multimodal associations across subsets of areas. Indeed, those target areas that are preferentially co-innervated by broadcasting neurons appear to have more similar visual response properties^26,27^. Broadcasting cells may also coordinate activity among the subset of areas they co-innervate, thus providing a signal which may link visual information across different processing streams. Furthermore, the highly divergent nature of signal transmission from a primary sensory cortex to its targets may help constrain models of hierarchical sensory processing. Beyond the functional implications, the mere existence of distinct projection motifs that either avoid or favor subsets of target areas raises the question of how these specific, long-range connectivity patterns are established during development.

## Methods

### Single neuron tracing

The anatomical single-cell tracing experiments were conducted at The Biozentrum, University of Basel, Switzerland. We licensed and performed all experimental procedures in accordance with institutional and cantonal animal welfare guidelines using both male and female adult (>8 weeks of age) C57BL/6 mice. Detailed protocols and all software are available at: *http://mouse.vision/han2017*

#### Two-photon guided single-cell electroporation

We performed surgery as described previously^32^. Briefly, we anesthetized animals with a mixture of fentanyl (0.05 mg kg^−1^), midazolam (5 mg kg^−1^) and medetomidine (0.5 mg kg^−1^), and maintained stable anaesthesia by isoflurane (0.5% in O_2_). We performed all electroporations on a custom linear scanning 2-photon microscope, equipped to image both a green and a red channel and running ScanImage 5.1^33^. For electroporation we used a patch pipette (12-16 MΩ) filled with plasmid DNA (pCAG-eGFP (Addgene accession 11150) or pAAV-EF1a-eGFP-WPRE (generous gift from Botond Roska; sequence file can be found in the Supplemental Materials, 100 ng/μl) and AlexaFluor 488 (50 μM) in intracellular solution, and delivered electroporation pulses (100 Hz, −14 V, 0.5 ms for 1 s) with an Axoporator 800A (Molecular Probes) when pushed against a target cell. We verified successful electroporation by dye filling of the cell body, and then sealed the skull with a chronic window using 1.5% agarose in HEPES-buffered artificial cerebrospinal fluid and a cover slip. We finally confirmed plasmid expression two days after electroporation by visualization of GFP epifluorescence through the chronic imaging window. Three to 10 days after electroporation, we transcardially perfused anesthetized mice with 10 ml 0.9% NaCl followed by 50 ml 4% paraformaldehyde in 0.1 M phosphate buffer (pH 7.4). We removed the brains from the skull and post-fixed them in 4% paraformaldehyde overnight. We then stored the fixed brains in PBS at 4 °C until imaging with serial-section 2-photon tomography.

#### Serial-section 2-photon tomography

We embedded the fixed brains in 5% oxidised agarose (derived from Sigma Type I agarose) and covalently cross-linked the brain to the agarose by incubation in an excess of 0.5–1% sodium borohydrate (NaBH4, Sigma) in 0.05 M sodium borate buffer overnight at 4°C. We then imaged embedded brains using a TissueVision 2-photon scanning microscope ^28^, which cut physical sections of the entire brain every 50 μm coronally, and acquired optical sections every 10 μm in two channels (green channel: 500-560 nm; red channel: 560-650 nm) using 940 nm excitation laser light (Mai Tai eHP, Spectraphysics). Each imaged section is formed from overlapping 800x800 μm “tiles”. We imaged with a resolution of 1 μm in x and y and measured an axial point spread function of ~5 μm FWHM using ScanImage 5.1.

#### Image processing and cell tracing

We stitched raw image tiles using a custom MATLAB-based software, *StitchIt*. *StitchIt* applies illumination correction based on the average tiles for each channel and optical plane, and subsequently stitches the illumination-corrected tiles from the entire brain. We then navigated through the stitched brain space using *MaSIV* (https://github.com/alexanderbrown/masiv), a MATLAB-based viewer for very large 3-D images, and traced axons using a custom, manual neurite-tracer extension for *MaSIV*.

To assign each voxel of the imaged brains to a brain area, we segmented each brain using areas defined by the Allen Reference Atlas (ARA, Common Coordinate Framework v3), after smoothing with a single pass of an SD=0.5 voxel Gaussian kernel using the Nifty “seg-maths” tool as described previously^34^. Briefly, we downsampled one imaging channel to a voxel size of 25 μm and converted it to MHD format using *StitchIt*. We then registered the volume to the ARA average template brain using Elastix^35^ by applying rigid affine transformation followed by non-rigid deformation with parameters as described previously^36,37^. We examined registration quality using a custom Python/PyQt5 application, *Lasagna,* which overlays the Allen template brain and the registered sample brain and is extendable to allow the overlay of traced cells, or the overlay of ARA area borders onto a down-sampled brain. In order to transform the traced cells into ARA space (sample to ARA) we calculated the inverse transform to the one calculated by Elastix (ARA to sample) and applied this to the traced points.

#### Analysis of traced neurons

To avoid potential incomplete filling of neurons from biasing the results of our analyses, we excluded cells with non-arborizing primary branches in the ipsilateral hemisphere from the analysis. Out of a total of 71 traced cells, we excluded 28 cells that exhibited abrupt, non-callosal terminations, thus restricting our analysis to ipsilateral projection patterns of 31 cells in V1 and 7 in other higher visual areas. Moreover, axonal branches terminating contralaterally or after entering the corpus callosum were considered as callosal terminations and were included in the analysis (compare ref^14^). We calculate the first order projection statistics only using the ARA-registered cells that satisfied these criteria. To reduce any artifacts associated with ARA registration or individual brain variability in boundaries between brain areas, we excluded any axon within 50 μm from any brain area boundary from the analysis. We then calculated the projection strength of each neuron to each area as the total length of axon of that neuron in an area. To determine the number of projection targets for every cell, we used a minimum projection strength of 1 mm axon length per target area.

### MAPseq

Animal procedures were approved by the Cold Spring Harbor Laboratory Animal Care and Use Committee and carried out in accordance with National Institutes of Health standards. Custom MATLAB (Mathworks) code for the analysis of projection patterns is available at: *http://mouse.vision/han2017.*

#### MAPseq sample processing

To define the V1 injection site and target higher visual areas LI, LM, AL, PM, AM and RL, we used optical imaging of intrinsic signals as previously described^26,38^. Briefly, we first implanted a customized head plate and then thinned the skull to increase its transparency. After 2-3 days of recovery, we sedated the mice (chlorprothixene, 0.7 mg/kg) and lightly anesthetized them with isoflurane (0.5-1.5% in O_2_), delivered via a nose cone. We illuminated visual cortex with 700 nm light split from an LED source into 2 light guides, performing imaging with a tandem lens macroscope focused 250-500 μm below the cortical surface and a bandpass filter centered at 700 nm with 10 nm bandwidth (67905; Edmund optics). We acquired images at 6.25 Hz with a 12- bit CCD camera (1300QF; VDS Vosskühler), frame grabber (PCI-1422; National Instruments) and custom software written in LabVIEW (National Instruments). We visually stimulated the contralateral eye of mice with a monitor placed at a distance of 21 cm and presented 25-35° patches of 100% contrast square wave gratings with a temporal frequency of 4 Hz and a spatial frequency of 0.02 cycles per degree for 2 s followed by 5 s of grey screen (mean luminance of 46 cd/m^2^). To establish a coarse retinotopy of the targeted area, we alternated the position of the patches: we used two different elevations (approx. 0 and 20°) and two different azimuths (approx. 60 and 90°); at each position we acquired at least 17 trials. We obtained intrinsic signal maps by averaging the responses during the stimulation time using ImageJ (National Institute of Mental Health, NIH) and mapping the location of the estimated spots of activation onto a previously acquired blood vessel picture.

We then pressure injected (Picospritzer III, Parker) 100 nl of 1x10^10^ GC/ml barcoded MAPseq Sindbis virus^23^ with a diversity of >8x10^6^ different barcode sequences unilaterally at a depth of 100-200 μm from the brain surface into V1 of a total of four 8-10 week old C57BL/6 females. In addition, we labeled the six higher visual areas by placing a DiI-coated micropipette into retinotopically matched positions according to intrinsic signal maps. For this, we allowed 2-5 μl of a 2.5 mg/ml DiI (Invitrogen D3911) in ethanol solution to dry on the outside of a pulled micropipette tip until some DiI crystals were visible. Mice were sacrificed 44-48 hours post-injection by decapitation, and their brain immediately extracted and flash frozen on dry ice.

We cut 180 μm thick coronal sections using a cryostat at −10°C blade and sample holder temperature, and melted each slice onto a clean microscope slide before rapidly freezing it on dry ice again. We then dissected each target area and the injection site using cold scalpels while keeping the brain sections frozen on a metal block cooled to approximately −20°C in a freezing 2.25M CaCl_2_ bath^39^. During dissection, we identified each dissected area using a fluorescent dissection microscope to visualize viral GFP expression and DiI stabs labeling each target area (Extended Data Fig. 7). Throughout the procedure, we took care to avoid sample cross-contamination by never reusing tools or blades applied to different areas and changing gloves between samples. To measure noise introduced by contamination, we collected samples of the olfactory bulb from each brain, which served as a negative control.

We then processed the dissected samples for sequencing largely as previously described^23^, but pooling all samples after first strand cDNA synthesis. Briefly, we extracted total RNA from each sample using Trizol reagent (Thermo Fisher) according to the manufacturer’s instructions. We mixed the sample RNA with spike-in RNA (obtained by *in vitro* transcription of a double stranded ultramer with sequence 5’-GTC ATG ATC ATA ATA CGA CTC ACT ATA GGG GAC GAG CTG TAC AAG TAA ACG CGT AAT GAT ACG GCG ACC ACC GAG ATC TAC ACT CTT TCC CTA CAC GAC GCT CTT CCG ATC TNN NNN NNN NNN NNN NNN NNN NNN NAT CAG TCA TCG GAG CGG CCG CTA CCT AAT TGC CGT CGT GAG GTA CGA CCA CCG CTA GCT GTA CA-3’ (IDT)^23^) and reverse transcribed the RNA mixture using gene specific primer 5’-CTT GGC ACC CGA GAA TTC CAN NNN NNN NNN NNX XXX XXX XTG TAC AGC TAG CGG TGG TCG-3’, where X_8_ is one of >300 trueseq like sample specific identifiers and N12 is the unique molecular identifier, and SuperscriptIV reverse transcriptase (Thermo Fisher) according to the manufacturer’s instructions. We then pooled all first strand cDNAs, purified them using SPRI beads (Beckman Coulter) and produced double stranded cDNA as previously described^40^. We then treated the samples using ExonucleaseI (NEB) and performed two rounds of nested PCR using primers 5’-CTC GGC ATG GAC GAG CTG TA-3’ and 5’-CAA GCA GAA GAC GGC ATA CGA GAT CGT GAT GTG ACT GGA GTT CCT TGG CAC CC GAG AAT TCC A-3’ for the first PCR and primers 5’-AAT GAT ACG GCG ACC ACC GA-3’ and 5’- CAA GCA GAA GAC GGC ATA CGA-3’ for the second PCR using Accuprime Pfx polymerase (Thermo Fisher). Finally, we gel extracted the resulting PCR amplicons using Qiagen MinElute Gel extraction kit according to the manufacturer’s instructions and sequenced the library on a Illumina NextSeq500 high-output run at paired-end 36 using the SBS3T sequencing primer for paired-end 1 and the Illumina small RNA sequencing primer 2 for paired-end 2.

#### MAPseq data analysis

Based on the sequencing results, we constructed a barcode matrix M of (number of barcodes) x (number of dissected areas) with entry *M_i,j_* representing the absolute counts of barcode *i* in area *j* as previously described^23^. We de-multiplexed the sequencing results, extracted the absolute counts of each barcode in each sample based on the UMI sequence and error corrected the barcode sequences, before matching barcode sequences to the virus library and constructing matrix M by matching barcode sequences across areas. We then filtered the barcode matrix for ‘high-confidence’ cell bodies inside the dissected area of V1 by requiring a minimum of 10 counts in at least one target area, an at least 10-fold difference between the cell body location in V1 and the most abundant target area in data normalized to the efficiency of library production as measured by the amount of recovered spike-in RNA counts, and an absolute minimum barcode count of 300 in V1. We then normalized the raw barcode counts in each area by the relative spike-in RNA recovery to the olfactory bulb sample, merged the results from all four processed brains into a single barcode matrix and used this matrix for all further analysis.

To determine whether a particular neuron projected to any given target area, we chose a conservative threshold of at least 5 barcode counts, based on the highest level of barcode expression in the olfactory bulb negative control sample.

#### Calculation of statistical significance of projection motifs

To calculate the statistical significance of broadcasting projection motifs, we compared against the simplest model in which we assumed that each neuron projected to each area independently. To generate predictions of this model, we first estimated the probability of projecting to each area, assuming independent projections. We define the probability *P*(*A_i_*) that a given neuron projects to the *i^th^* area *A_i_* as

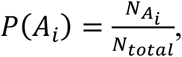

where 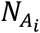 is the number of neurons in the sample that project to area *A_i_*, *i* = 1.. *k* for k analyzed target areas, and *N_total_* is the total number of neurons in the sample.

In our MAPseq experiments, we do not have direct access to *N_total_*, since for technical reasons we only include neurons that have at least one projection among the dissected targets. Since in principle some neurons might project to none of the areas dissected (see Fig. 3a), failure to include these would lead to an underestimate of *N_total_*. However, assuming independence of projections we can infer *N_total_* from the available measurements.

To estimate *N_total_*, we first observe that

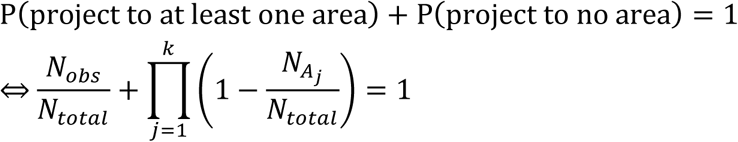

where *N_obs_* is the total number of neurons observed to project to at least one area. For *k=6* areas, we can expand this expression to

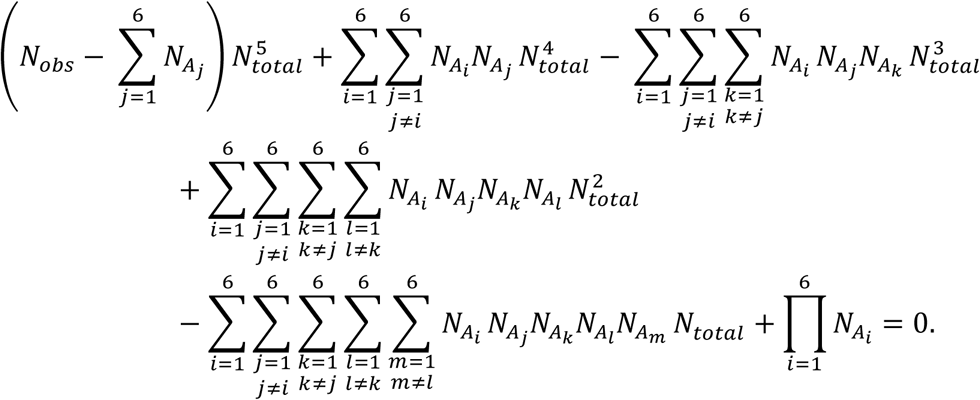

Noting that this is a quintic equation in *N_total_*, we can use a root finder to solve for *N_total_* numerically, and use the result to calculate *P*(*A_i_*).

Using the derived *N_total_* and *P*(*A_i_*), we can calculate the p-value for every possible broadcasting motif by calculating the value of the binomial cumulative distribution function, for a total of *N_total_* tries, the empirical number of observed counts (successes), and P(motif) assuming independent projections. We calculated the p-value of all possible bi-, tri- and quadfurcations, and determined significantly over- or under-represented broadcasting motifs at a significance threshold of α=0.05 after Bonferoni correction.

### Data availability

All sequencing data are publicly accessible on the Sequence Read Archive under accessions SRR5274845 (ZL097 for mouse 4 and mouse 5) and SRR5274844 (ZL102 for mouse 6 and mouse 7). All single cell tracing results are accessible on http://mouse.vision/han2017 and will be uploaded to http://neuromorpho.org.

## Acknowledgements

We would like to thank Ashley Juavinett, Longwen Huang, Sonja Hofer and Petr Znamenskiy for comments on the manuscript. Funding sources: National Institutes of Health (5RO1NS073129 to A.M.Z., 5RO1DA036913 to A.M.Z.); Brain Research Foundation (BRF-SIA-2014-03 to A.M.Z.); IARPA (MICrONS to A.M.Z.); Simons Foundation (382793/SIMONS to A.M.Z.); Paul Allen Distinguished Investigator Award (to A.M.Z.); PhD fellowship from the Boehringer Ingelheim Fonds (to J.M.K.); PhD fellowship from the Genentech Foundation (to J.M.K); European Research Council (NeuroV1sion 616509 to T.D.M.-F.), and Swiss National Science Foundation (SNSF 31003A_169802 to T.D.M.-F.).

## Author contributions

Y.H. generated the dataset for fluorescence-based axonal tracing. D.C. and Y.H. traced the cells. R.A.A.C. analyzed the serial 2-P imaging data and axonal projection patterns. J.M.K. and F.I. collected the MAPseq dataset. J.M.K. and A.M.Z. performed the analysis of projection patterns. J.M.K., T.D.M-F. and A.M.Z. wrote the paper.

## Author information

The authors declare no conflict of interests.

## Supplemental Notes

Like any other method, MAPseq is subject to false positives (detection of an extra, artefactual projection) and false negatives (failing to identify a real connection). Please refer to ref^23^, for a detailed discussion of the effect of fibers of passage, co-infections, infection of more than one cell with the same barcode sequence, and various other sources of false negatives and false positives. Below, we briefly discuss the most important considerations.

### Supplemental Note 1: Unique labeling of cells by viral infection

As described in more detail previously^23^, in MAPseq we deliver barcodes to cells by random viral infection. In the simplest scenario we aim to deliver one unique barcode per labeled cell, such that each cell can unequivocally be identified by a single sequence. In practice however, there are two scenarios that deviate from this simple model.

On the one hand, we might infect cells with more than one virus particle and thus label each cell with more than one barcode sequence. Such multiple labeling results in MAPseq overestimating the total number of traced cells, but does not result in a false projection pattern and maintains the relative proportions of projection types. Such multiple labeling will therefore not lead to any false positive results.

On the other hand, degenerate labeling, that is labeling more than one cell with the same barcode sequence, produces artificial projection patterns that result from the merged projection pattern of all the cells labeled with the same barcode. We avoid degenerate labeling in MAPseq by infecting cells with a very large virus library. The fraction of uniquely labeled cells for a given size of virus library with an empirically determined barcode probability distribution can be described by

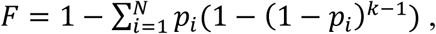

where *p_i_* is the probability of barcode *i*= 1..*N* to be chosen from the virus library, *k* is the number of labeled cells and *N* is the total number of barcodes in the library^23^. Given the size of the library used in this study (~10^7^ distinct barcode sequences) and the number of cells traced per brain (~300), the vast majority of cells (>99 %) will be uniquely labeled. In previous work^23^ we validated these theoretical predictions by multiple independent experimental methods.

### Supplemental Note 2: Other sources of false positives and negatives

Beyond errors introduced by degenerate labeling (see above), MAPseq is subject to false positives and negatives from other sources. False negatives are introduced into the dataset when the strength of a real projection falls below the detection floor of MAPseq. Conversely, MAPseq false positives are introduced when barcodes are detected in areas in which they were not originally present in (e.g. by sample cross-contamination).

Several lines of evidence suggest that MAPseq false negative and positive rates are low. In previous experiments ^23^, we measured the efficiency of MAPseq to be very similar to that of Lumafluor retrobeads (91.4 ± 6 % (mean ± std error))^23^, and therefore concluded that MAPseq false negative rates are comparable to those of other, well established methods. Similarly, we found MAPseq false positive rates to be low (1.4 ± 0.8 % (mean ± std error))^23^.

In the present study we improve on these previous estimates by comparing MAPseq data directly to the gold standard of single neuron tracing. To do so, we first used a bootstrapping procedure to measure the minimum pairwise cosine distances between each member of a randomly sampled set of MAPseq neurons and the remaining MAPseq neurons. We then measured the minimum pair-wise cosine distance between the fluorescence-based single neuron reconstructions and the remaining MAPseq neurons. As a negative control, we further measured the minimum pairwise distance between a set of random neurons (with their projection strengths drawn from a uniform distribution) and the remaining MAPseq neurons. We then compared the distribution of minimum distances for the three sets of measurement and found that while the MAPseq-to-MAPseq and fluorescence-based reconstructions-to-MAPseq distributions are statistically indistinguishable (two sample Kolmogorov-Smirnov test, p=0.9439), both the MAPseq-to-MAPseq and fluorescence-based reconstructions-to-MAPseq distributions are statistically different from the random neuron-to-MAPseq distribution (Fig. 2d; two sample Kolmogorov-Smirnov test, p=2.75x10^-4^ and p=8.76x10^-5^, respectively). Taken together, these results indicate that the statistics of projections inferred by MAPseq are indistinguishable from those obtained by fluorescence-based single neuron reconstructions.

## Extended Data Figures

**Extended Data Figure 1:**
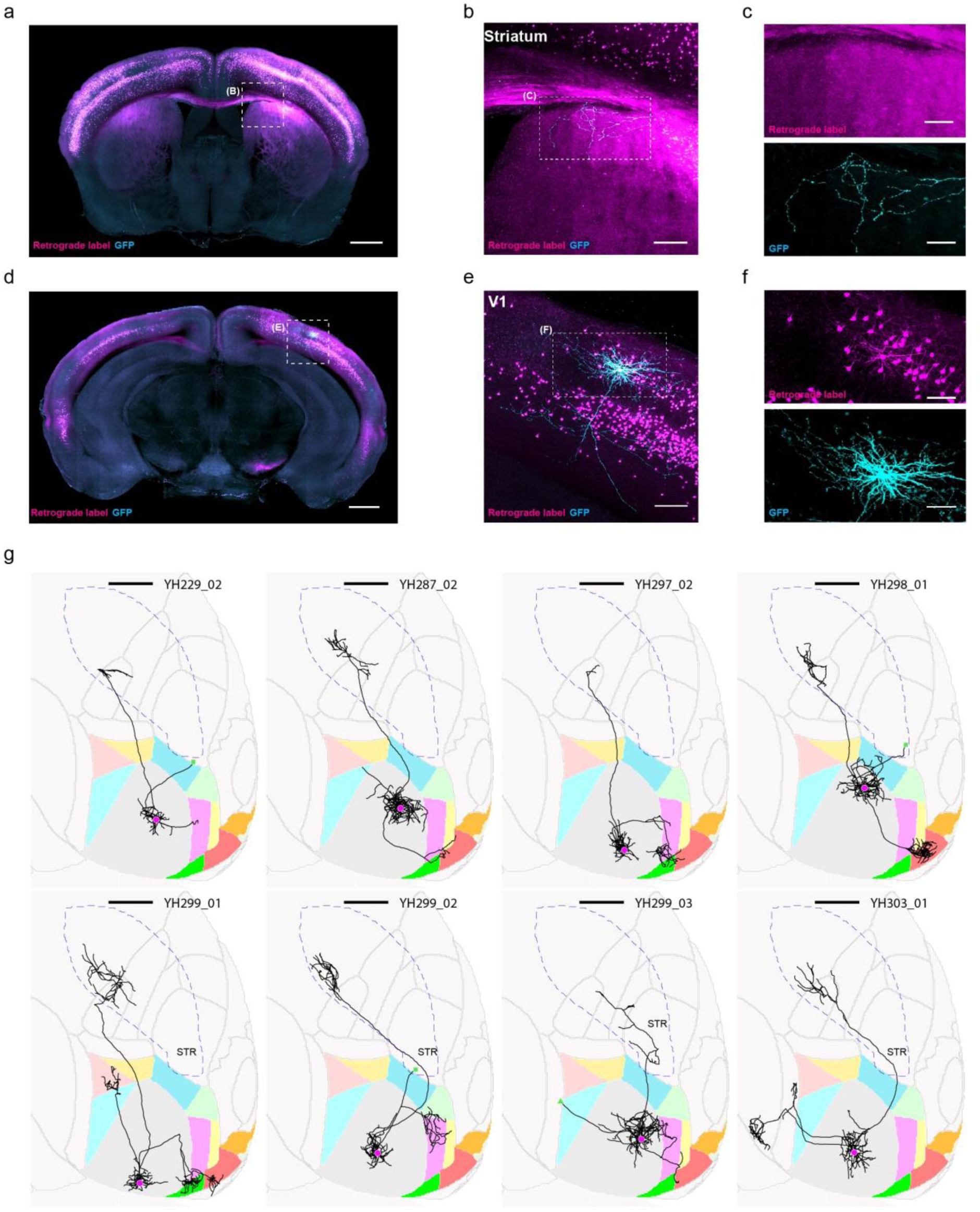
Single-neuron tracing protocol efficiently fills axons projecting to the ipsilateral striatum. We retrogradely labeled striatum projecting cells by stereotactically injecting cholera toxin subunit B conjugated with AlexaFluro594 or PRV-cre into the visual striatum of wild type mice or tdTomato reporter mice (Ai14, JAX), respectively (magenta). With visual guidance of two-photon microscopy, we electroporated single retrogradely labeled cells in V1 with a GFP expressing plasmid (cyan). (**a**) Coronal, maximum intensity projections of visual striatum. Scale bar = 1 mm. (**b**) Higher magnification view of the visual stratum. Scale bar = 0.2 mm. (**c**) Single channel images of the same axonal arbor as in (**b**). (**d**) Coronal maximum intensity projection containing V1. Scale bar = 1 mm. (**e**) Higher magnification view of V1. Scale bar = 0.2 mm. (**f**) Single channel images of V1. Scale bar = 0.2 mm. (**g**) Horizontal ARA-space projections of eight retrogradely labeled and electroportated cells. Cell ID numbers are indicated at the top right of each thumbnail. Scale bar = 1 mm. Note that one additional cell was retrogradely labeled and electroporated, which revealed its axonal projection to the striatum, but it is not shown because the brain was too distorted to allow accurate atlas registration.

**Extended Data Figure 2:**
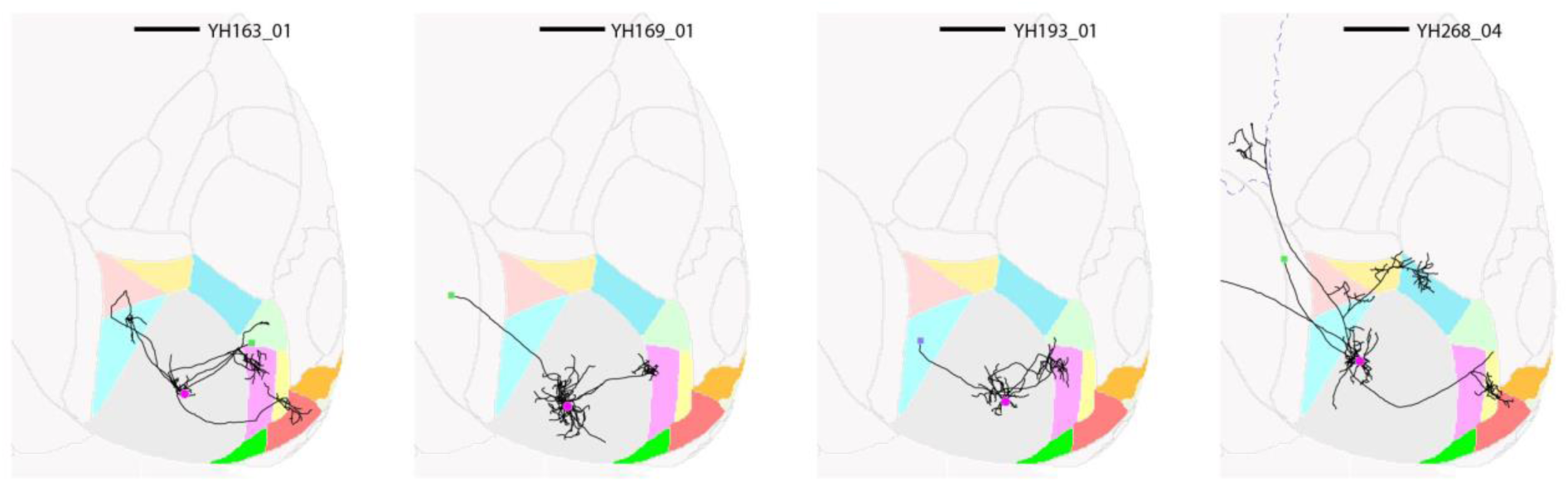
Some axonal branches terminate abruptly without arborizing, while other branches of the same neuron arborise extensively within different target areas and appear to be completely filled. Abrupt terminations in white matter are labeled with a green square. Abrupt terminations in grey matter are labeled with a blue square. Horizontal views of the ARA space are shown, and cell ID numbers are indicated at the top right of each thumbnail. Scale bar = 1 mm.

**Extended Data Figure 3:**
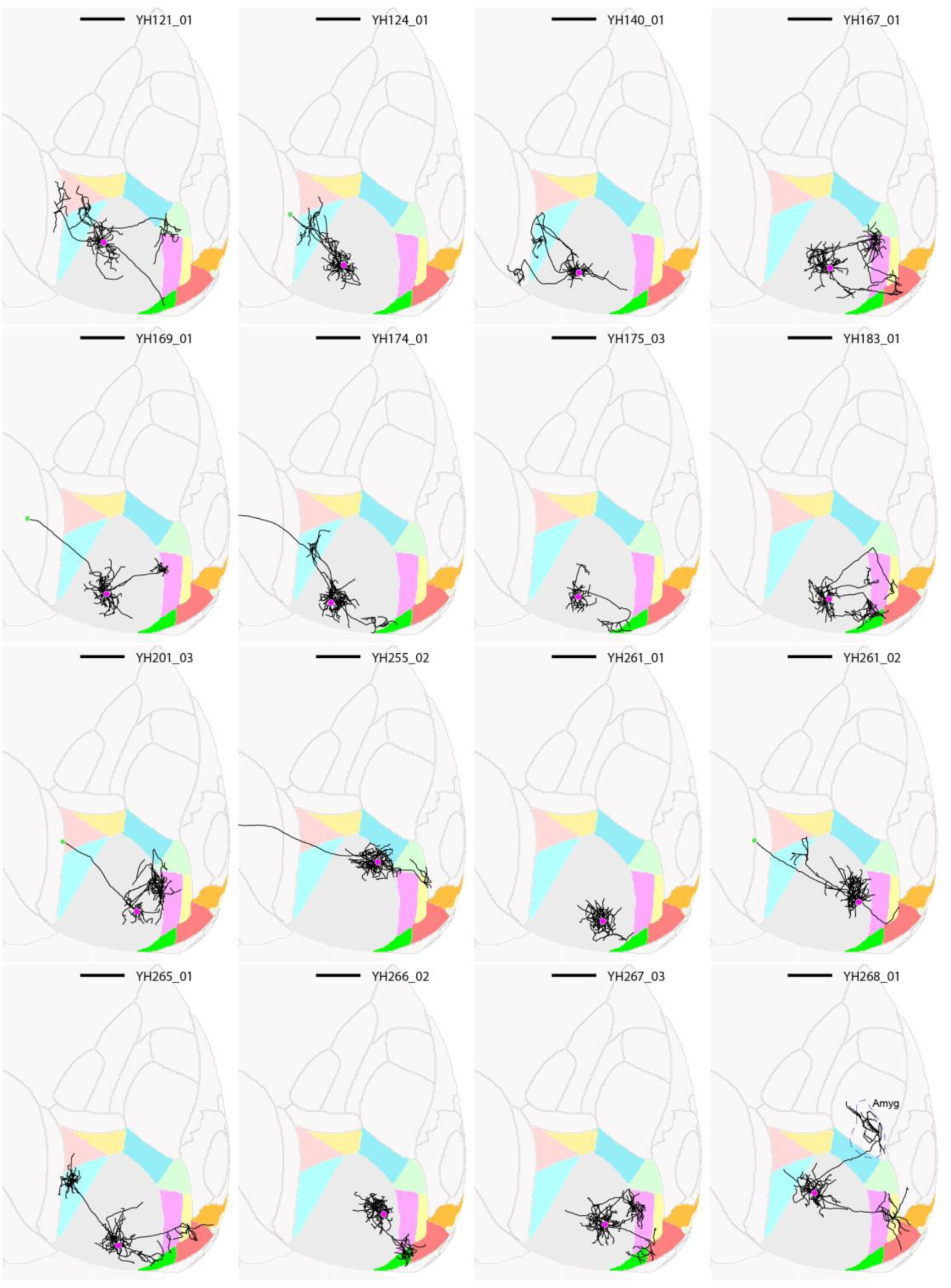
Thumbnails of traced layer 2/3 V1 neurons, part 1. Horizontal views of the ARA space are shown, and cell ID numbers are indicated at the top right of each thumbnail. Scale bar = 1 mm.

**Extended Data Figure 4:**
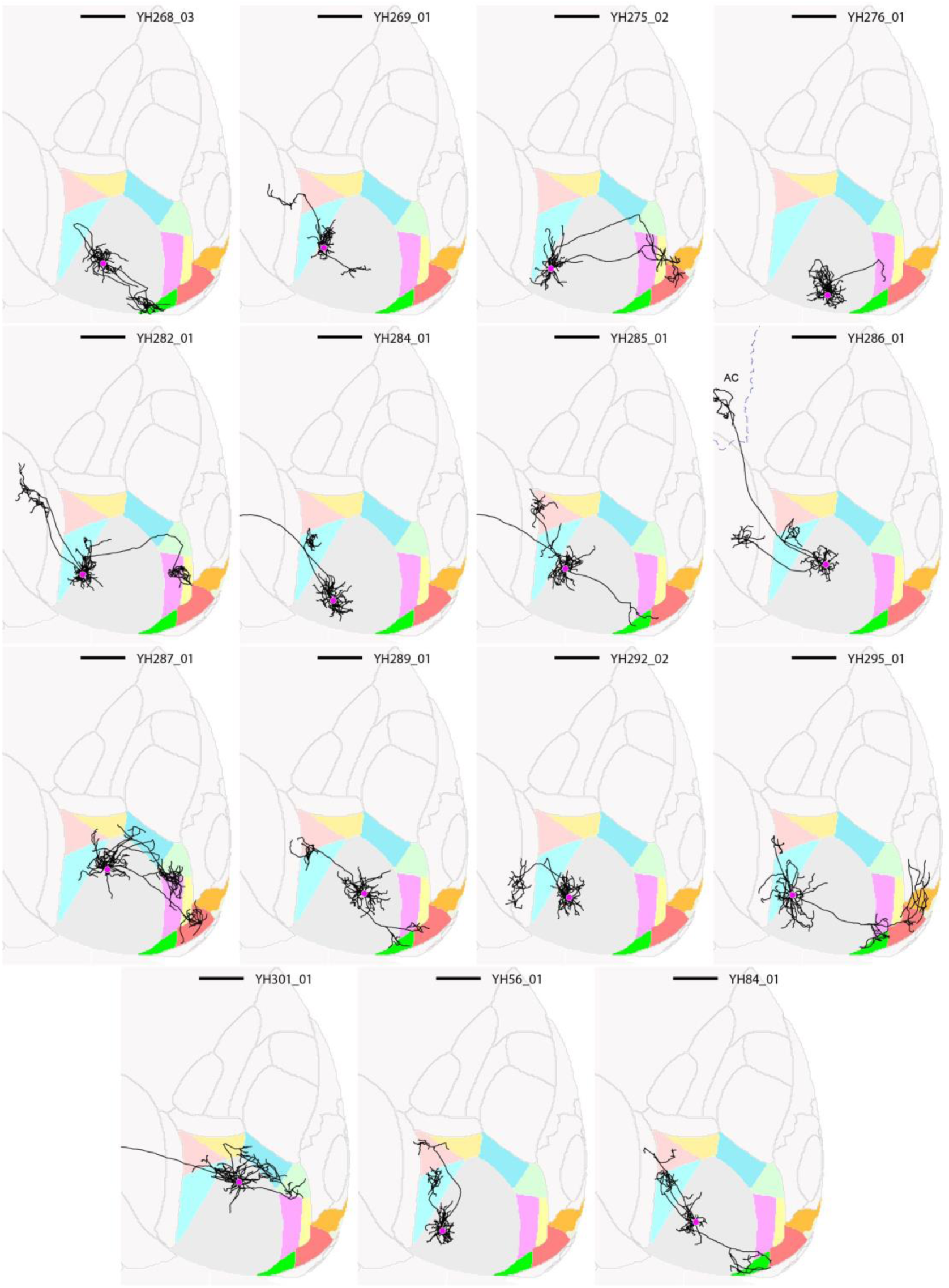
Thumbnails of traced layer 2/3 V1 neurons, part 2. Horizontal views of the ARA space are shown, and cell ID numbers are indicated at the top right of each thumbnail. Scale bar = 1 mm.

**Extended Data Figure 5:**
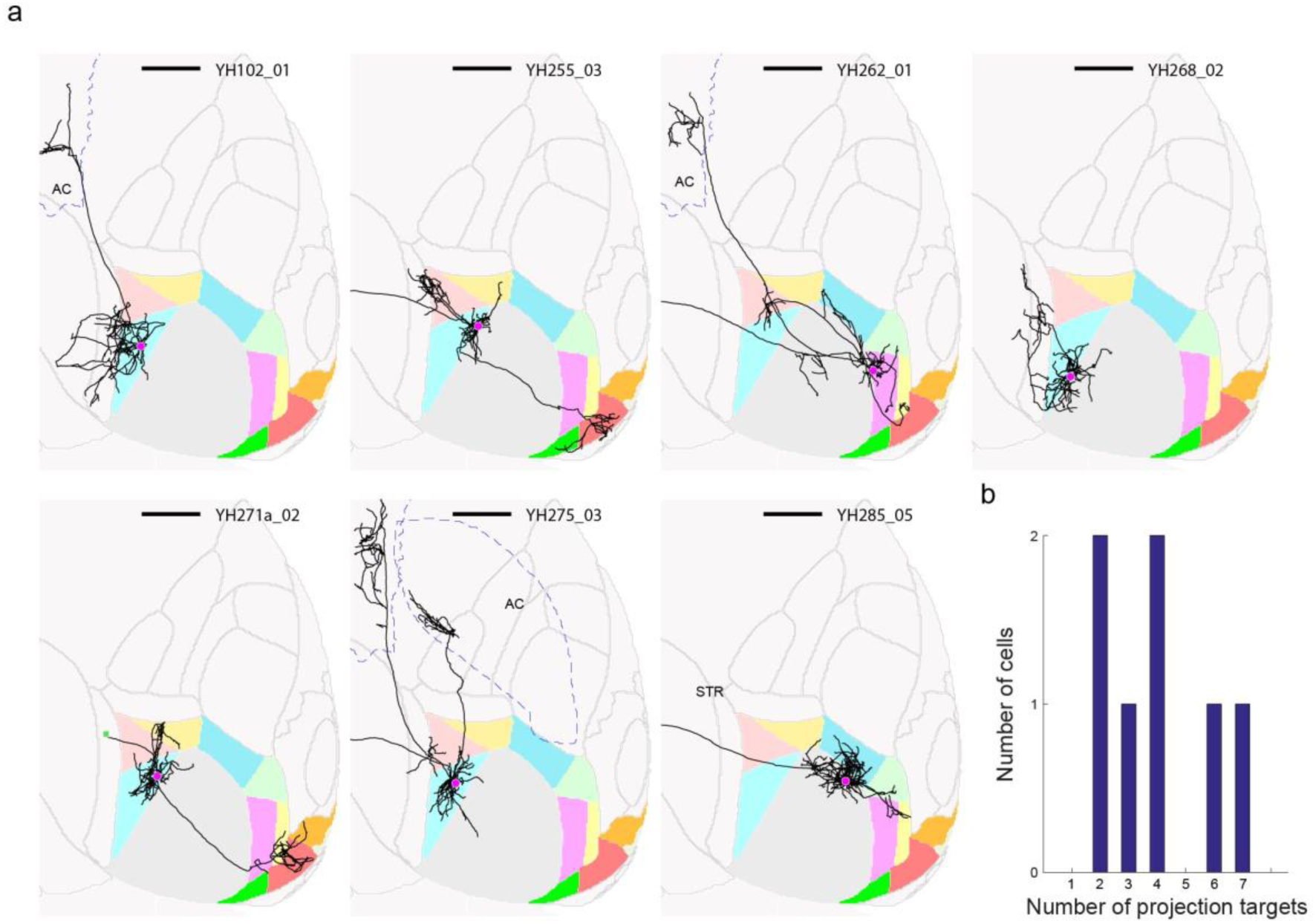
Individual neurons in higher visual areas project to more than one target area. (**a**) Thumbnails of all traced neurons with cell bodies not in V1. Brain area identity is color-coded as in Figure 1. Cell identity is indicated at the top right of each thumbnail. Scale bar = 1 mm. (**b**) Histogram of the number of target areas per cell.

**Extended Data Figure 6:**
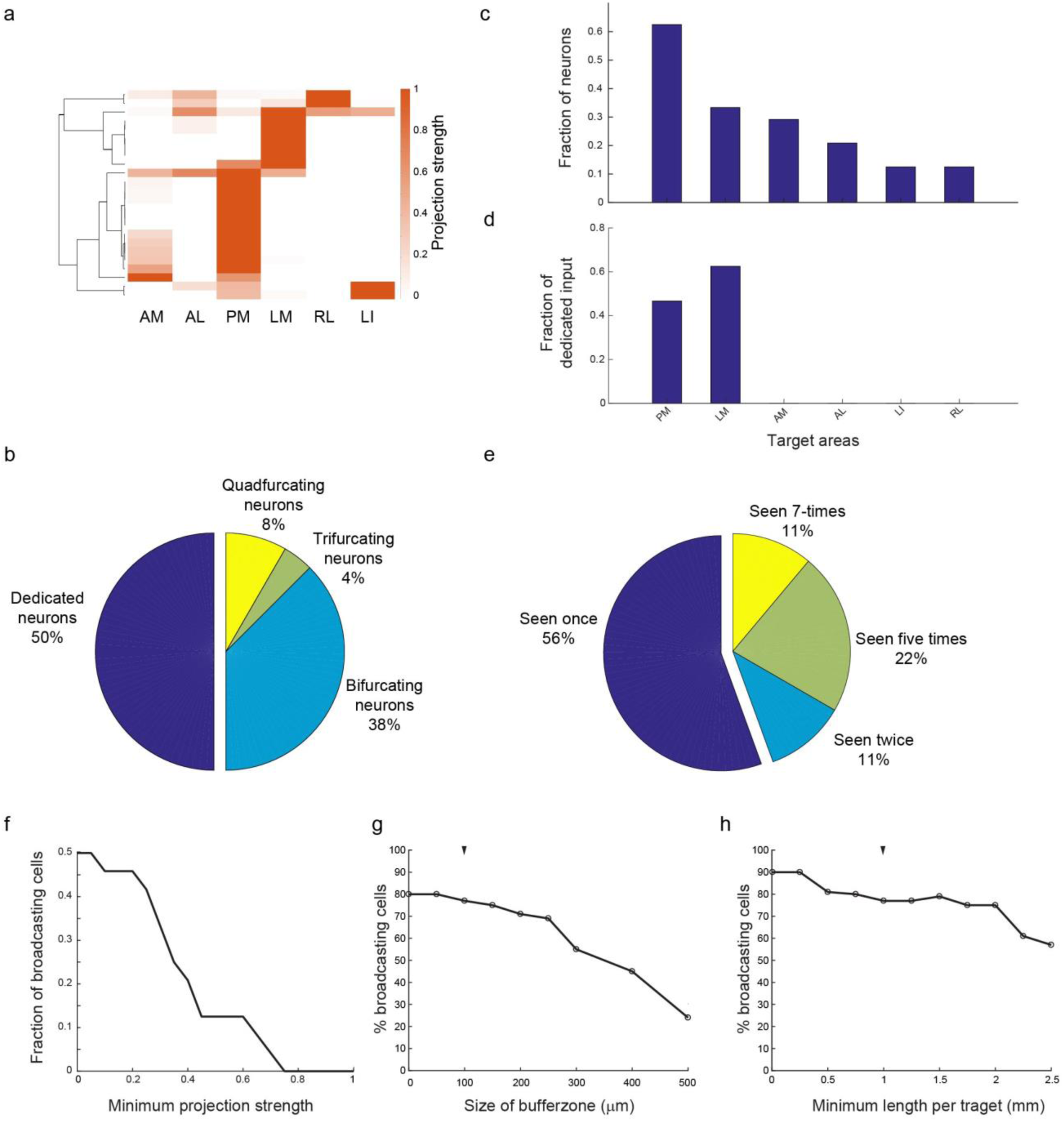
Conclusions from fluorescence-based single neuron tracing data hold true if analysis is restricted to subset of target areas. (**a**) The projection patterns of reconstructed GFP-filled neurons when only the six target areas LI, LM, AL, PM, AM, and RL are considered. Projection strengths are normalized to the maximum projection of each neuron, and only neurons projecting to at least one target area are shown. (**b**) Pie chart showing the distribution of target area numbers per projecting neuron. (**c**) Bar graph illustrating the fraction of all cells projecting to each target area. (**d**) The fraction dedicated input per area. (**e**) The number of times each binarized projection motif is observed. (**f**) The fraction of broadcasting cells as a function of the minimum projection strength (relative to the primary target) that each area needs to receive to be considered a target. (**g**) The fraction of broadcasting cells as a function of increasing buffer zones between areas within which axons are ignored, assuming a minimum projection of 1 mm of axon per target area. (**h**) The fraction of broadcasting cells as a function of the minimal amount of axon per area for it to be considered a target, assuming buffer zones of 100 μm width.

**Extended Data Figure 7:**
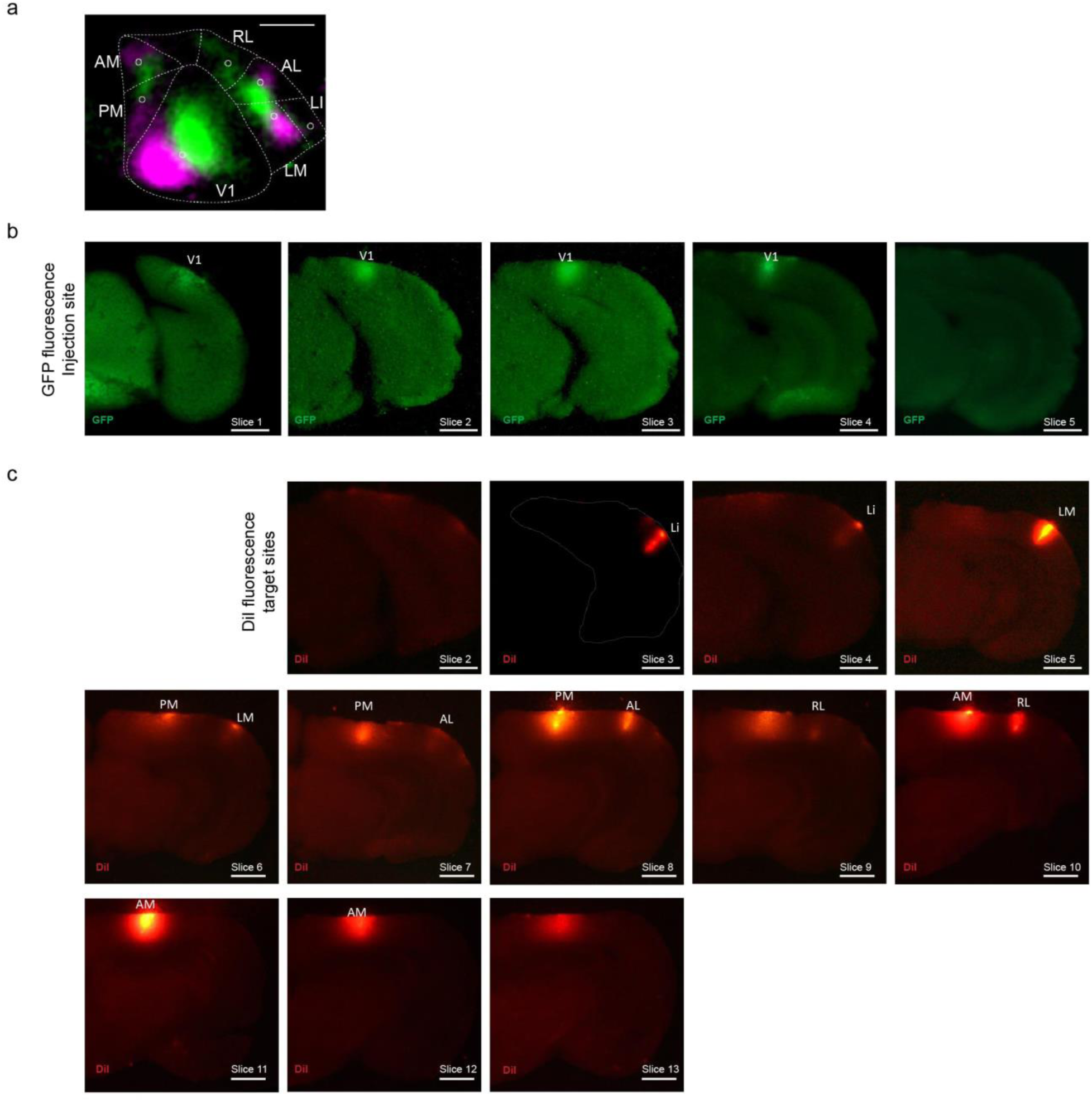
MAPseq dissection strategy. We identified the to-be-dissected higher visual areas by performing intrinsic imaging of visual cortex in response to stimuli at different positions in the contralateral visual field and mapping the resulting changes in intrinsic signals. (**a**) A representative retinotopic map, with responses to the two 25° visual stimuli pseudocolored in green and magenta (stimulus 1 position: 90° azimuth, 20° elevation; stimulus 2 position: 60° azimuth, 20° elevation). Based on this map, we fluorescently labelled retinotopically matched positions in the to-be-dissected cortical areas with a DiI stab (white circles). Putative borders between the higher visual areas are indicated in dashed lines for orientation. Scale bar = 1 mm. (**b**) The MAPseq virus injection site is discernible in consecutive frozen 180 μm thick coronal sections, using GFP fluorescence. Scale bar = 1 mm. (**c**) DiI injections targeted to matched retinotopic positions in six target areas identified by intrinsic signal imaging. DiI epifluorescence images of each 180 μm thick slice are shown, and dissected areas are labeled. Scale bar = 1 mm.

**Extended Data Figure 8:**
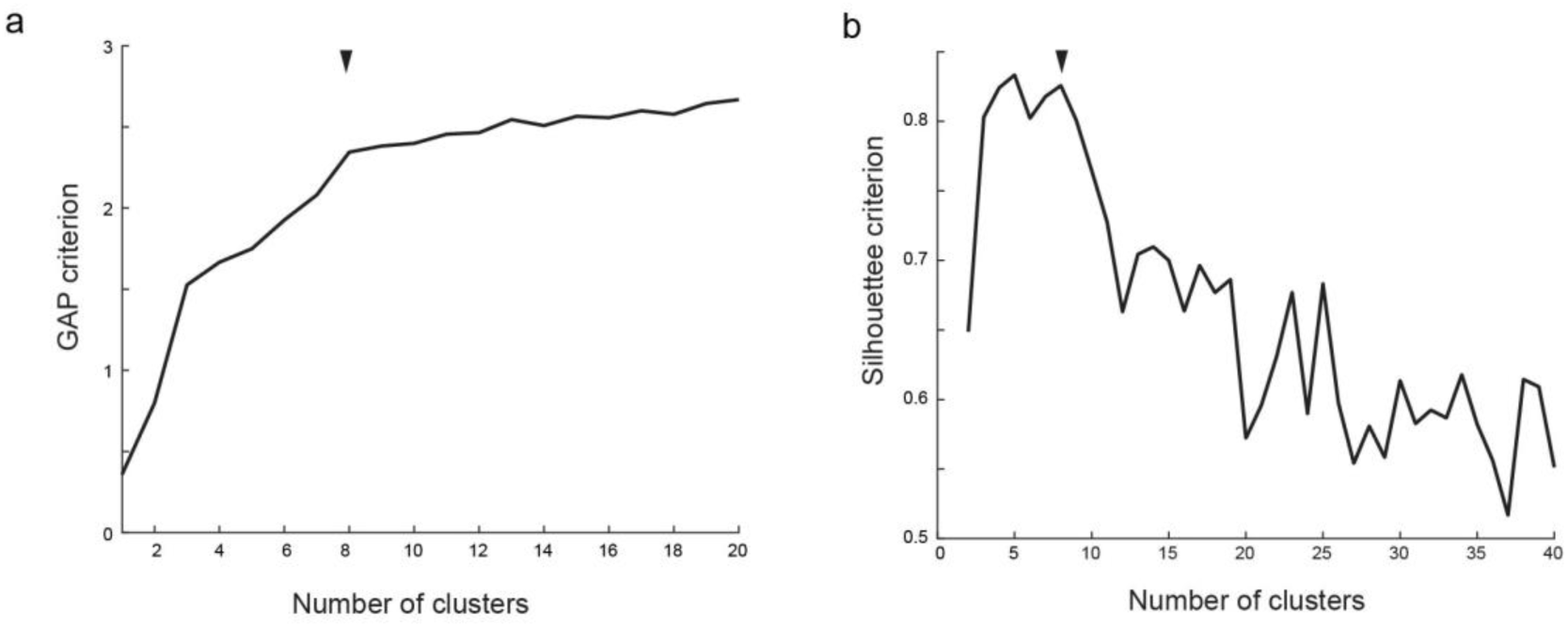
Clustering of MAPseq data. (**a**) GAP and (**b**) Silhouette criteria for k-means clustering of the MAPseq neurons as a function of the number of clusters. Black arrow heads indicate chosen number of clusters (k=8).

## Extended Data Tables

**Extended Data Table 1:**
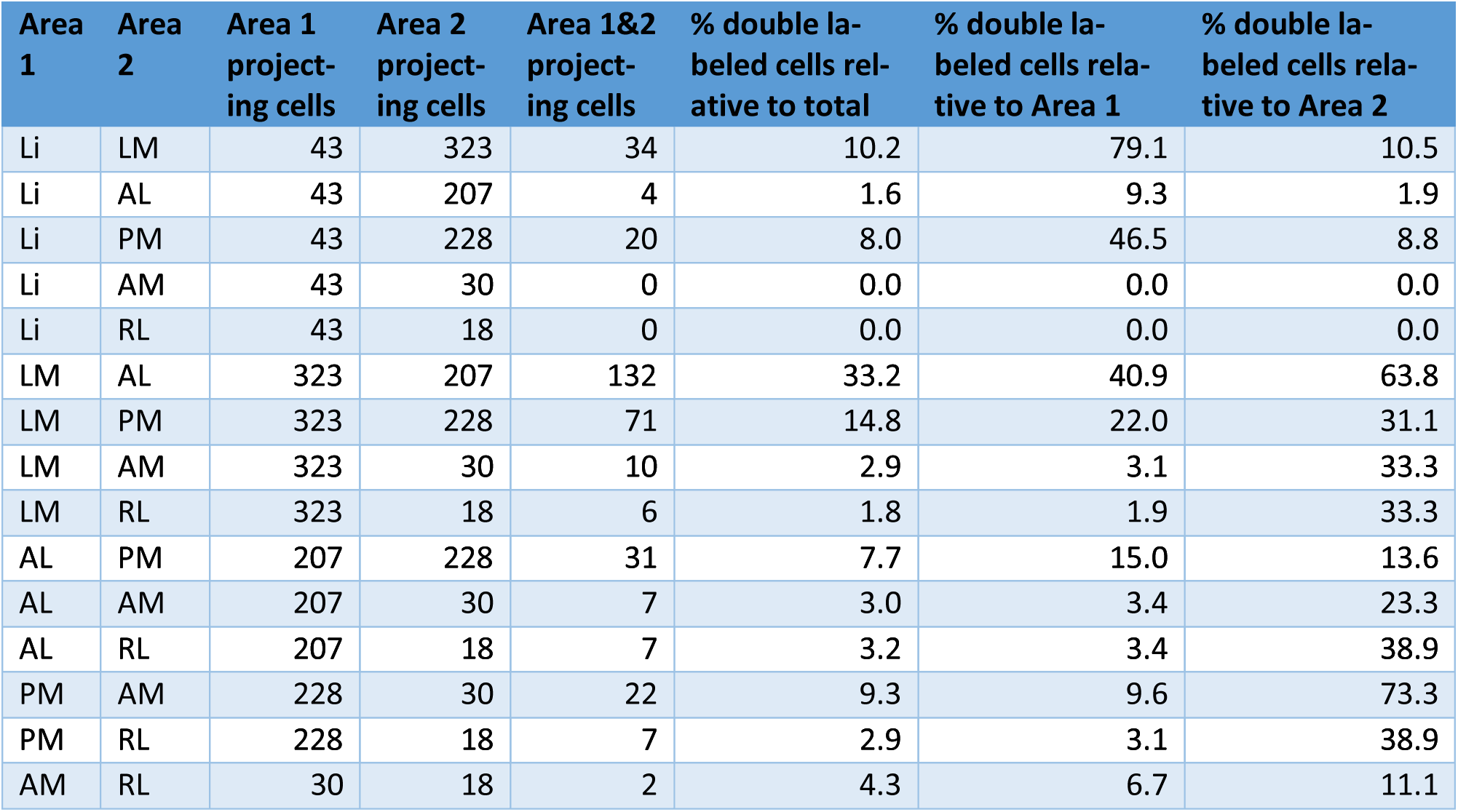
Simulated double retrograde tracing based on MAPseq anterograde tracing data. We determined whether any given MAPseq neuron targeted any one area using the same projection criterion used for the analysis in Fig. 3. For any pair of the six higher visual areas analyzed using MAPseq, we then determined the number of cells that projected to either area in the pair, or to both — effectively simulating double retrograde tracing from the two areas in the pair to V1. We here show the raw counts of cell projecting to each area, and the percentages of cells that project to the indicated pairs of areas, i.e. “double labeled” cells.

## References

1. Zingg, B. et al. Neural networks of the mouse neocortex. Cell 156, 1096–1111 (2014).

2. Oh, S. W. et al. A mesoscale connectome of the mouse brain. Nature 508, 207–214 (2014).

3. Markov, N. T. et al. Cortical High-Density Counterstream Architectures. Science 342, 1238406–1238406 (2013).

4. Felleman, D. J. & Van Essen, D. C. Distributed hierarchical processing in the primate cerebral cortex. Cereb. Cortex 1, 1–47 (1991).

5. Rockland, K. S. Collateral branching of long-distance cortical projections in monkey. J. Comp. Neurol. 521, 4112–4123 (2013).

6. Wang, Q. & Burkhalter, A. Area map of mouse visual cortex. J. Comp. Neurol. 502, 339–357 (2007).

7. Nakamura, H., Gattass, R., Desimone, R. & Ungerleider, L. The modular organization of projections from areas V1 and V2 to areas V4 and TEO in macaques. J. Neurosci. 13, (1993).

8. Segraves, M. & Innocenti, G. Comparison of the distributions of ipsilaterally and contralaterally projecting corticocortical neurons in cat visual cortex using two fluorescent tracers. J. Neurosci. 5, (1985).

9. Sincich, L. C. & Horton, J. C. Independent Projection Streams from Macaque Striate Cortex to the Second Visual Area and Middle Temporal Area. J. Neurosci. 23, 5684–5692 (2003).

10. Glickfeld, L. L., Andermann, M. L., Bonin, V. & Reid, R. C. Cortico-cortical projections in mouse visual cortex are functionally target specific. Nat. Neurosci. 16, 219–226 (2013).

11. Sato, T. R. & Svoboda, K. The Functional Properties of Barrel Cortex Neurons Projecting to the Primary Motor Cortex. J. Neurosci. 30, 4256–4260 (2010).

12. Chen, J. L., Carta, S., Soldado-Magraner, J., Schneider, B. L. & Helmchen, F. Behaviour-dependent recruitment of long-range projection neurons in somatosensory cortex. Nature 499, 336–340 (2013).

13. Yamashita, T. & Petersen, C. C. H. Target-specific membrane potential dynamics of neocortical projection neurons during goal-directed behavior. Elife 5, 2221 (2016).

14. Yamashita, T. et al. Membrane Potential Dynamics of Neocortical Projection Neurons Driving Target-Specific Signals. Neuron 80, 1477–1490 (2013).

15. Movshon, J. A. & Newsome, W. T. Visual response properties of striate cortical neurons projecting to area MT in macaque monkeys. J. Neurosci. 16, 7733–7741 (1996).

16. Nassi, J. J. & Callaway, E. M. Parallel processing strategies of the primate visual system. Nat. Rev. Neurosci. 10, 360–372 (2009).

17. Massé, I. O., Régnier, P. & Boire, D. in Axons and Brain Architecture 93–116 (2016). doi: 10.1016/B978-0-12-801393-9.00005-0

18. Bullier, J. & Kennedy, H. Axonal bifurcation in the visual system. Trends Neurosci. 10, 205–210 (1987).

19. Weisenhorn, D. M. V., Ilung, R. B. & Spatz, W. B. Morphology and connections of neurons in area 17 projecting to the extrastriate areas mt and 19DM and to the superior colliculus in the monkey Callithrix jacchus. J. Comp. Neurol. 362, 233–255 (1995).

20. Ding, S.-L., Van Hoesen, G. & Rockland, K. S. Inferior parietal lobule projections to the presubiculum and neighboring ventromedial temporal cortical areas. J. Comp. Neurol. 425, 510–530 (2000).

21. Economo, M. N. et al. A platform for brain-wide imaging and reconstruction of individual neurons. Elife 5, 13 (2016).

22. Gong, H. et al. High-throughput dual-colour precision imaging for brain-wide connectome with cytoarchitectonic landmarks at the cellular level. Nat. Commun. 7, 12142 (2016).

23. Kebschull, J. M. et al. High-Throughput Mapping of Single-Neuron Projections by Sequencing of Barcoded RNA. Neuron 91, 975–987 (2016).

24. Hartline, H. K. The response of single optic nerve fibers of the vertebrate eye to illumination of the retina. Am. J. Physiol. (1938).

25. Lehky, S. R. & Sejnowski, T. J. Network model of shape-from-shading: neural function arises from both receptive and projective fields. Nature 333, 452–4 (1988).

26. Andermann, M. L., Kerlin, A. M., Roumis, D. K., Glickfeld, L. L. & Reid, R. C. Functional specialization of mouse higher visual cortical areas. Neuron 72, 1025–1039 (2011).

27. Marshel, J. H. H., Garrett, M. E. E., Nauhaus, I. & Callaway, E. M. M. Functional Specialization of Seven Mouse Visual Cortical Areas. Neuron 72, 1040–1054 (2011).

28. Ragan, T. et al. Serial two-photon tomography for automated ex vivo mouse brain imaging. Nat. Methods 9, 255–258 (2012).

29. Osten, P. & Margrie, T. W. Mapping brain circuitry with a light microscope. Nat. Methods 10, 515–523 (2013).

30. Wang, Q., Sporns, O. & Burkhalter, A. Network Analysis of Corticocortical Connections Reveals Ventral and Dorsal Processing Streams in Mouse Visual Cortex. J. Neurosci. 32, (2012).

31. Smith, I. T., Townsend, L. B., Huh, R., Zhu, H. & Smith, S. L. Stream-dependent development of higher visual cortical areas. Nat. Neurosci. 20, 200–208 (2017).

32. Pecka, M., Han, Y., Sader, E. & Mrsic-Flogel, T. D. Experience-Dependent Specialization of Receptive Field Surround for Selective Coding of Natural Scenes. Neuron 84, 457–469 (2014).

33. Pologruto, T. A., Sabatini, B. L. & Svoboda, K. ScanImage: flexible software for operating laser scanning microscopes. Biomed. Eng. Online 2, 13 (2003).

34. Niedworok, C. J. et al. aMAP is a validated pipeline for registration and segmentation of high-resolution mouse brain data. Nat. Commun. 7, 711879 (2016).

35. Klein, S., Staring, M., Murphy, K., Viergever, M. A. & Pluim, J. elastix: A Toolbox for Intensity-Based Medical Image Registration. IEEE Trans. Med. Imaging 29, 196–205 (2009).

36. Kim, Y. et al. Whole-Brain Mapping of Neuronal Activity in the Learned Helplessness Model of Depression. Front. Neural Circuits 10, 3 (2016).

37. Renier, N. et al. Mapping of Brain Activity by Automated Volume Analysis of Immediate Early Genes. Cell 165, 1789–1802 (2016).

38. Roth, M. M. et al. Thalamic nuclei convey diverse contextual information to layer 1 of visual cortex. Nat. Neurosci. 19, 299–307 (2015).

39. Bryan, W. P. & Byrne, R. H. A calcium chloride solution, dry-ice, low temperature bath. J. Chem. Educ. 47, 361 (1970).

40. Morris, J., Singh, J. M. & Eberwine, J. H. Transcriptome Analysis of Single Cells. JoVE e2634--e2634 (2011). doi:10.3791/2634

